# Platelet proteo-transcriptomic profiling validates mediators of thrombosis and proteostasis in patients with myeloproliferative neoplasms

**DOI:** 10.1101/2023.10.23.563619

**Authors:** Sarah Kelliher, Sara Gamba, Luisa Weiss, Zhu Shen, Marina Marchetti, Francesca Schieppati, Caitriona Scaife, Stephen Madden, Kathleen Bennett, Anne Fortune, Su Maung, Michael Fay, Fionnuala Ní Áinle, Patricia Maguire, Anna Falanga, Barry Kevane, Anandi Krishnan

## Abstract

Patients with chronic Myeloproliferative Neoplasms (MPN) including polycythemia vera (PV) and essential thrombocythemia (ET) exhibit unique clinical features, such as a tendency toward thrombosis and hemorrhage, and risk of disease progression to secondary bone marrow fibrosis and/or acute leukemia. Although an increase in blood cell lineage counts (quantitative features) contribute to these morbid sequelae, the significant qualitative abnormalities of myeloid cells that contribute to vascular risk are not well understood. Here, we address this critical knowledge gap via a comprehensive and untargeted profiling of the platelet proteome in a large (n= 140) cohort of patients (from two independent sites) with an established diagnosis of PV and ET (and complement prior work on the MPN platelet transcriptome from a third site). We discover distinct MPN platelet protein expression and confirm key molecular impairments associated with proteostasis and thrombosis mechanisms of potential relevance to MPN pathology. Specifically, we validate expression of high-priority candidate markers from the platelet transcriptome at the platelet proteome (*e.g.,* calreticulin (CALR), Fc gamma receptor (FcγRIIA) and galectin-1 (LGALS1) pointing to their likely significance in the proinflammatory, prothrombotic and profibrotic phenotypes in patients with MPN. Together, our proteo-transcriptomic study identifies the peripherally-derived platelet molecular profile as a potential window into MPN pathophysiology and demonstrates the value of integrative multi-omic approaches in gaining a better understanding of the complex molecular dynamics of disease.

**Highlights:** MPN patient platelet proteome identifies key pathobiological mediators of thrombosis and proteostasis. The MPN platelet proteomic profile validates our prior findings from the platelet transcriptome.

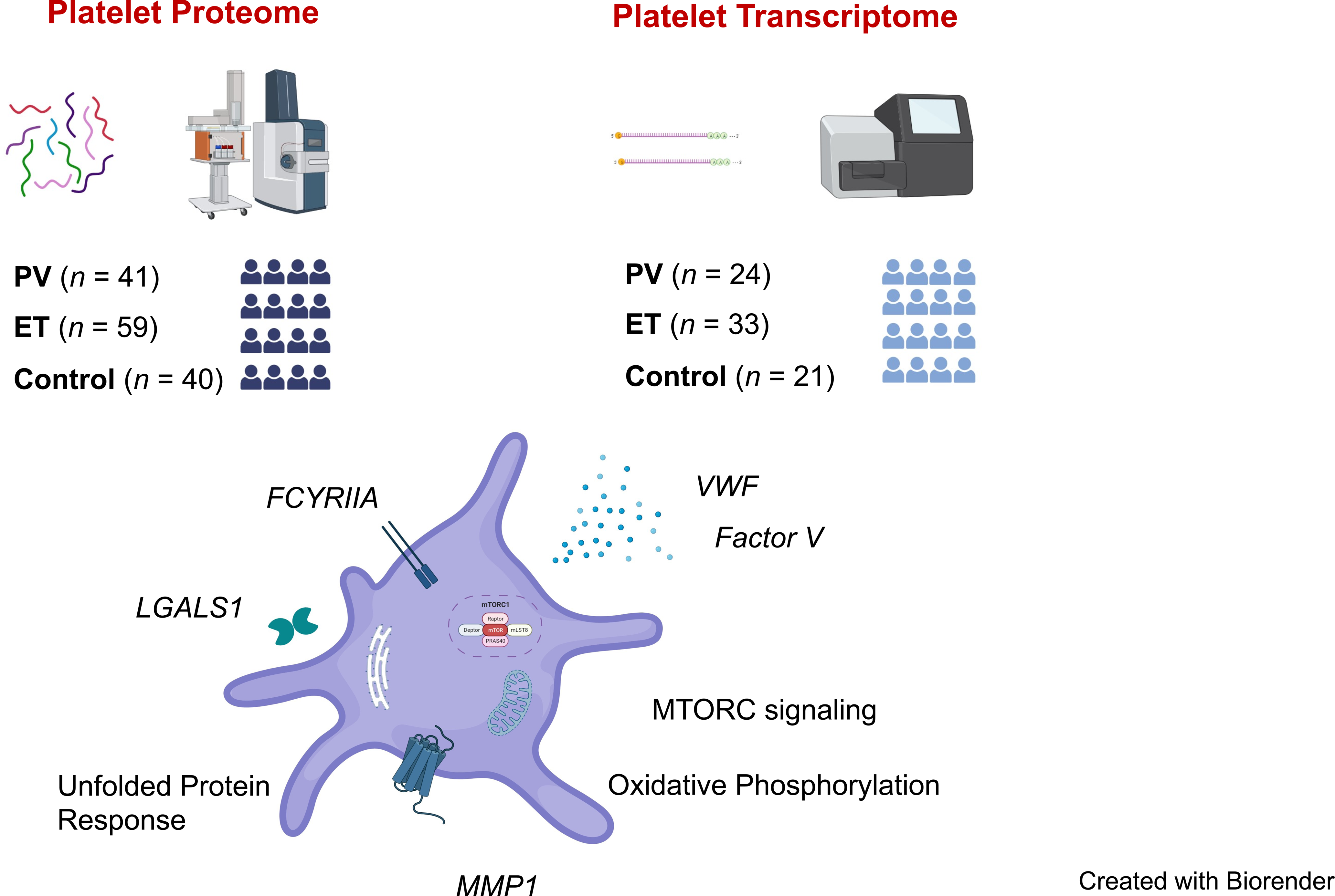

## Introduction

Myeloproliferative neoplasms (MPN) are chronic bone marrow malignancies characterised by clonal proliferation of hematopoietic precursors and elevated cell counts in peripheral blood. The three most common Philadelphia-chromosome negative (Ph^-^) MPN subtypes polycythemia vera (PV), essential thrombocythemia (ET) and myelofibrosis (MF), are diagnosed according to morphologic, clinical, laboratory, and genetic criteria^1,2^. Patients with MPN are at risk of progression to secondary bone marrow fibrosis or acute leukemia and experience a substantial burden of microvascular symptoms, negatively impacting their quality of life^3-5^. However, it is thrombosis, including both arterial and venous events, which represents the leading cause of morbidity and mortality for patients with PV and ET^6-9^. Several patient and disease-specific characteristics have been shown to correlate with increased clinical risk, yet despite currently available standard of care, the prediction and prevention of disease complications remains challenging^10^.

The discovery of the *JAK2* V617F mutation in 2005 heralded a new era in the management of MPN, and the application of genomic techniques has radically modified diagnostic and prognostic approaches in the interim^11^. In addition to the MPN driver mutations (*JAK2* V617F, *CALR*, *MPL* & *JAK2* exon 12) there is increasing interest in other somatic gene mutations including epigenetic regulators, spliceosome components, transcription factors and oncogenes^12^. These parameters are key informants of risk stratification for patients with MF (MIPSS70^13^), yet clinical utility lags behind for the more indolent subtypes (PV & ET) where additional genomic information currently has limited impact on patient management^14^. Thus, greater understanding of fundamental MPN pathobiology, in particular, the molecular and cellular basis of disease, is urgently required to discover novel mediators that can be leveraged for prognostic and therapeutic purposes in these chronic MPN subtypes.

Platelets have a well-established and integral role in maintaining vascular integrity and mediating hemostasis. Additionally, platelets and their cargo have been shown in numerous translational studies to influence a myriad of pathological processes including inflammation, immune responses, and malignancy^15-17^. Platelet hyperactivity and the procoagulant role of platelets is a hallmark of MPNs^18^ ^19,20^. Yet, a comprehensive picture of the MPN platelet molecular profile is still lacking. In recent published work that serves as a prelude to this data^21^, we evaluated a large (n=120) cohort of MPN patients of all three chronic subtypes, and discovered progressive changes in platelet transcriptomic expression thus offering novel candidate signatures and deeper insights into MPN pathobiology (*e.g.* interferon response, immuno-thrombosis, and impaired proteostasis). However, to date, no studies have evaluated the unbiased platelet proteome in a sizeable clinical cohort of patients with MPN and the overlap of MPN platelet proteo-transcriptome therefore, not understood.

In the present study, with the objective of delineating the MPN proteo-transcriptomic landscape, we performed a comprehensive and untargeted quantitative profiling of the platelet proteome in a large (n= 140) cohort of patients with an established diagnosis of PV and ET. We demonstrate marked differences in platelet protein expression and discover novel MPN-altered pathways in platelets. Importantly, we validate, across two independent sites, candidate signatures from the MPN platelet transcriptome at the protein level. Together, this amounts to a comprehensive molecular profile of differential MPN pathobiology at high-sensitivity cellular resolution.

## Results

### Platelet proteo-transcriptome across multi-site MPN clinical cohorts

Patient samples were recruited for untargeted, quantitative proteomic analysis. Using established and standardized platelet isolation protocols (see Methods), we prepared purified platelets from peripheral blood samples of patients with an established diagnosis of MPN and a cohort of healthy controls recruited across two sites: Hospital Papa Giovanni XXIII, Bergamo, Italy and Mater Misericordiae University Hospital, Dublin, Ireland. Together, this platelet proteome compendium comprised of 140 human peripheral blood samples from healthy donors (n= 40) and World Health Organization-defined MPN patients (n= 59 ET, n= 41 PV). Pertinent clinical features are listed in **Table 1A** and shown in **Figure 1**; including MPN subtype (**A**), somatic driver mutation status (**B**), sex (**C**), treatment (**D**), age (**E**), and platelet count (**F**). Any inter-patient variability in patient age, sex, and treatment as well as experimental batch effects were adjusted as confounding factors in all downstream expression analyses (see Methods).

**Figure 1:**
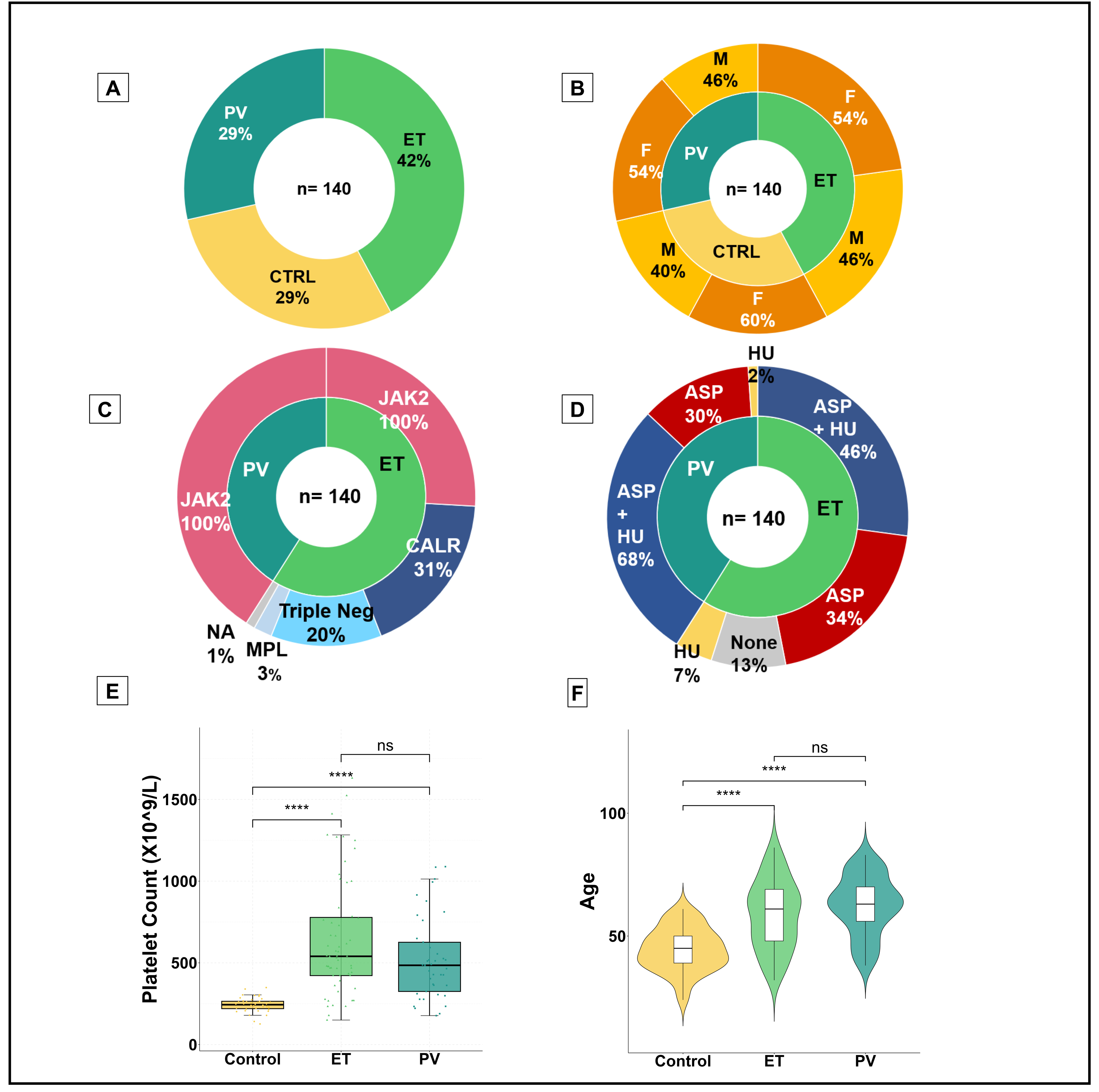
Patient variables representing MPN clinical cohort for proteomic work package. **A,** Similarity in distribution of MPN subtypes and controls, with slightly higher proportion ET (PV n= 41, ET n= 59, control n= 40) **B**, Comparable and balanced distribution of sex across MPN subtypes and controls. Larger percentage of female healthy controls. **C,** All patients with PV harbored the *JAK2* V617F mutation, and in keeping with the general ET population *JAK2* V617F was the most common driver mutation followed by *CALR* and *MPL*, with 12 triple negative ET patients included in this study**. D,** MPN patient therapies reflecting current clinical practice. The majority of PV and ET patients were prescribed aspirin (ASA), with hydroxyurea (HU) as a commonly utilized cytoreductive therapy. To control for any inter-patient variability, all treatment, in addition to patient, sex and experimental batch are adjusted as confounding factors in downstream differential expression analyses **E,** Comparable distribution of age across MPN subtypes and controls. Violin plots of patient age from each MPN subtype reflect clinical expectation, with slightly higher median age noted for ET and PV patients compared to controls. **F,** Platelet counts, as box plots, measured at the same date and time as experimental platelet sampling. As expected, Mann-Whitney *U* tests marked by asterisks indicate a statistically significant difference between control and MPN groups (*** *p*= <0.001, ns = not significant).

**Table 1A:**
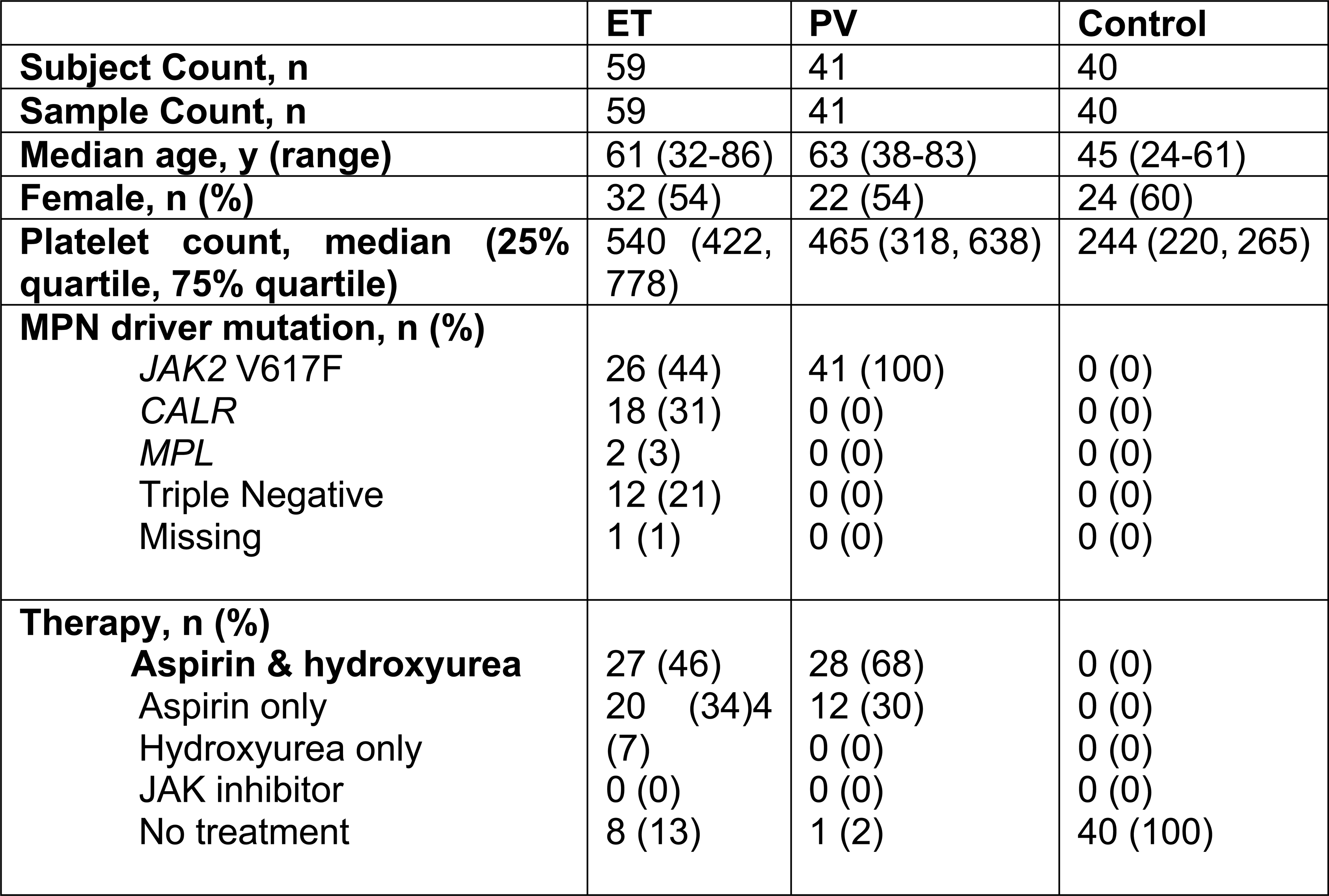
Characteristics of Patients & Controls Included in the Proteomic Cohort. MPN patient characteristics recruited across two sites (Papa Giovanni XXIII Hospital, Bergamo, Italy and Mater Misericordiae University Hospital, Dublin, Ireland) for proteomic analysis.

For comparative proteo-transcriptomic analysis, we leveraged our published dataset of the MPN platelet transcriptome comprising of 120 human peripheral blood samples from healthy donors (n= 21) and World Health Organization–defined MPN patients (n= 99) (single center, Stanford University, CA). Clinical variables of patients from this independent cohort (PV n= 33; ET n= 24, control n= 21) are outlined in **Table 1B** and described extensively in the original publication^21^.

**Table 1B:** Characteristics of Patients & Controls Included in the Transcriptomic Cohort. MPN patient characteristics recruited at single center (Stanford, USA^21^) included here for comparative transcriptomic analysis. Key demographic and other patient variables (listed below) have no statistically significant difference (unpaired t-test) between the (transcriptome) and Mater/Bergamo proteome cohorts. These variables include: age, sex, platelet count, and distribution of MPN driver mutation (*JAK2, CALR, MPL*) and therapy (aspirin, hydroxyurea).

|  | **ET** | **PV** | **Control** |
| --- | --- | --- | --- |
| **Subject Count, n** | 24 | 31 | 21 |
| **Sample Count, n** | 24 | 33 | 21 |
| **Median age, y (range)** | 58 (30-78) | 60 (27-84) | 57 (26-69) |
| **Female, n (%)** | 16 (66) | 14 (45) | 5 (24) |
| **Platelet count, median**<br>**(25% quartile, 75% quartile)** | 562<br>(464,799) | 447 (310, 557) | 167 (146,185) |
| **MPN driver mutation, n (%)** |  |  |  |
| *JAK2* V617F | 13 (54) | 31 (100) | 0 (0) |
| *CALR* | 7 (30) | 0 (0) | 0 (0) |
| *MPL* | 2 (8) | 0 (0) | 0 (0) |
| Triple Negative | 2 (8) | 0 (0) | 0 (0) |
| Missing | 0 (0) | 0 (0) | 0 (0) |
| **Therapy, n (%)** |  |  |  |
| **Aspirin& hydroxyurea** |  |  |  |
| Aspirin only | 6 (25) | 10 (32) | 0 (0) |
| Hydroxyurea only | 10 (42) | 12 (39) | 0 (0) |
| JAK inhibitor | 5 (21) | 1 (3) | 0 (0) |
| No treatment | 1 (4) | 7 (23) | 0 (0) |
|  | 2 (8) | 3 (9) | 21 (100) |

### MPN platelet proteome distinguishes disease phenotype and reveals prothrombotic, proinflammatory, and profibrotic signatures

Focusing for this study on the more prothrombotic subtypes of MPNs (PV and ET), we hypothesized that the platelet proteome differs in MPN subtypes and would offer insights into the underlying pathophysiology and the associated thrombotic phenotype. We compared platelet proteomic expression in PV and ET with that of healthy donors and discovered distinct signatures. Unsupervised principal component analysis (PCA) of MPN patients and controls (**Figure 2A**) confirmed that the collective variability from the first two principal components was MPN disease status (26% of total variance, after adjusting for age, sex, treatment, and experimental batch).

**Figure 2:**
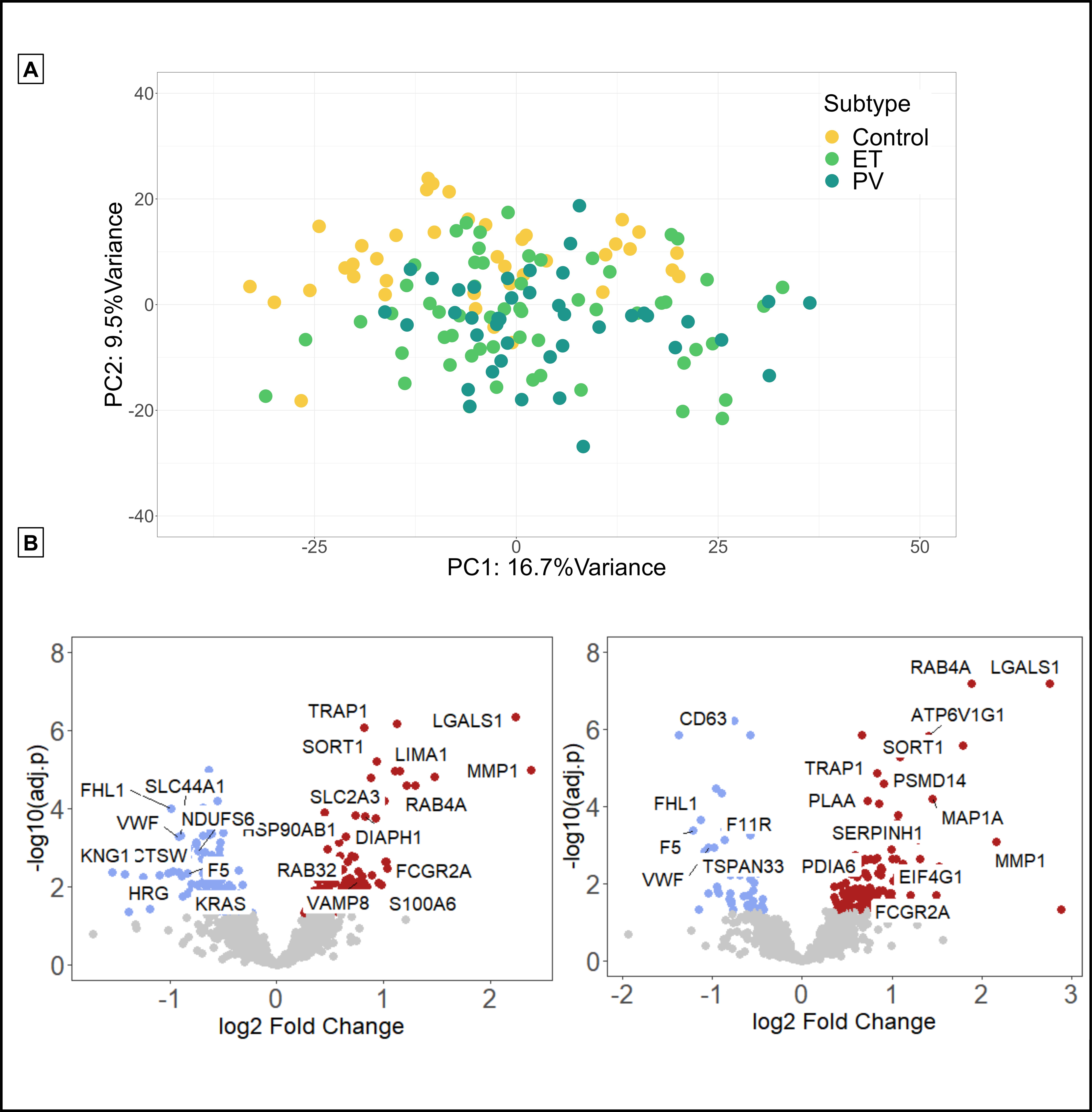
MPN platelet proteome distinguishes disease phenotype and reveals subtype specific signatures. **A,** Unsupervised principal component analysis (PCA) of normalized platelet protein expression adjusted for age, sex, treatment (antiplatelet and cytoreduction) and experimental batch. PC1 and PC2 colored by MPN subtype; and each contrasted with controls (n= 40, yellow): ET (n= 59, ET (n= 59, light green), PV (n= 41, dark green). The first two principal components account for 26% of total variance in the data. **B,** Volcano plots (two panels of ET, PV) of differential protein expression showing log2 fold change versus statistical significance (negative log10 of *p*-values) of each gene. Significant up-regulated and down-regulated genes are those with *p*-values (FDR) smaller or equal to 0.05 and absolute value of fold changes larger or equal to 1.5.

A total of 1,952 platelet proteins were quantified across MPN and control samples. Differential expression (DE) analysis of proteomic data (volcano plot, **Figure 2B, C**) efficiently distinguished each of the MPN subtypes and resulted in highly significant expression signatures (false discovery rate FDR <0.05) with 227 proteins differentially regulated in ET (113 increased and 114 decreased), and 166 in PV (122 increased and 44 decreased) compared to healthy donors.

Differential markers in ET and PV highlight candidate proteins as potential mediators of the clinically encountered pro-thrombotic phenotype, including proteins associated with hypercoagulation (e.g. MMP1, SERPINH1, FcγRIIA, PDIA6) and drivers of inflammation (LGALS1, S100A6, SLC25A24, CD63). Select candidates are highlighted in **Figure 2B, C** and discussed in detail in **Table 2**. Also, while the profibrotic phenotype of myelofibrosis is well-understood^22^, candidate markers (LGALS1, S100A6) in the early, more indolent subtypes of these chronic myeloid malignancies point to potential platelet mediators of fibrosis of relevance to MF.

**Table 2.** Description Select/representative candidate proteins that may variably influence the proinflammatory, pro-thrombotic, and profibrotic processes in MPNs.

| **Candidate protein**<br>**(Gene name)** | **Platelet differential**<br>**expression by MPN subtype**<br>**Absolute fold change**<br>**(direction of change)** | **Cellular function and relationship to procoagulant,**<br>**proinflammatory, and profibrotic pathways in other**<br>**published work.** |
| --- | --- | --- |
| **MMP1** | **PV:** 4.5 (increased)<br>**ET:** 5.2 (increased) | MMP1, an interstitial collagenase, is known to cleave protease-activated receptor 1 (PAR1) and promote platelet activation and regulate thrombogenesis in vitro<sup>69-71</sup>. Furthermore, MMP1 mediates tumor invasion by compromising vascular barrier integrity and has been associated with inferior prognosis in solid organ malignancies<sup>72-74</sup>. |
| **FcγRIIA** | **PV:** 2 (increased)<br>**ET:** 2 (increased) | Unbalanced FcγRIIA-mediated platelet aggregation was previously reported to promote thrombosis<sup>75</sup>. |
| **SERPINH1** | **PV:** 2 (increased) | There is evidence of decreased expression or ablation of this collagen binding, platelet adhesion protein in immobilized mammals as a thromboprotective mechanism<sup>76-78</sup>. |
| **LGALS1** | **PV:** 6.7 (increased)<br>**ET:** 4.7 (increased) | Galactin-1 is a beta-galactosidase binding protein which is reported to promote tumour cell proliferation and survival in haematological malignancies<sup>79</sup>. Recent data delineates the contribution of galactin-1 to disease severity in myelofibrosis and identified the protein as a potential drug target with disease modifying effects<sup>80</sup>. |
| **S100A6** | **ET:** 2 (increased) | The S100 family of proteins is a major player in hematopoietic proliferation and recent work has identified proinflammatory/profibrotic roles for S100A6, S100A8, S100A9 in bone marrow, granulocytes, and plasma in MPN<sup>80-85</sup>. |
| **PDIA6** | **PV:** 1.7 (increased) | Protein disulfide isomerases (PDIs) are key mediators of platelet ER homeostasis and the relationship between PDIs, ER & oxidative stress, platelet activation and thrombosis has recently been elucidated<sup>86,87</sup>. |
| **HPSE** | **PV:** 1.8 (increased) | Heparanase cleaves heparan sulfate proteoglycans and participates in extracellular matrix remodeling. It has been shown to be increased in the plasma extracellular vesicles of patients with PV<sup>56</sup>. |
| **DIAPH1** | **PV:** 1.9 (increased)<br>**ET:** 1.8 (increased) | We find increased expression of protein diaphanous homolog-1 (DIAPH1) possibly suggesting altered megakaryopoiesis in peripherally circulating platelets in PV and ET. DIAPH regulates proplatelet formation via Rho-mediated actin polymerization and microtubule assembly<sup>88</sup>. |
| **RAB4A** | **PV:** 3.7 (increased)<br>**ET:** 2.3 (increased) | A Ras GTPase signaling protein which regulates platelet alpha granule release<sup>89</sup>. |
| **CD63** | **PV: 2.6** (decreased) | Downregulation of CD63 has been associated with proliferation and metastasis in solid organ malignancy regulated by IL-6, IL-27 and STAT3 signaling<sup>90</sup>. |
| **CTSC** | **PV: 2** (increased) | Abundant Cathepsin C drives inflammation through macrophage activation via NF-κB signaling pathway<sup>91</sup>. |
| **VAMP8** | **ET: 1.7** (increased) | VAMP8 regulates platelet granule secretion and thrombosis *in vivo*<sup>92</sup>. |
| **EIF4G1** | **PV: 2.1** (increased)<br>**ET: 1.74** (increased) | EIF4G1 has been identified as a prognostic biomarker in breast cancer<sup>93</sup>. |
| **HSP90AB1** | **ET: 1.5** (increased) | We find increased expression of heat shock proteins in ET (HSP90AB1, TRAP1) and PV (HYOU1, HSPH1, DNAJA2). Heat shock protein 70 kDa and heat shock protein 90 kDa are two families of chaperone networks with integral roles in protein folding, degradation, trafficking, and maturation. They are known to promote oncogenesis by protecting a spectrum of cancer related proteins<sup>94-96</sup>. |
| **SLC25A2** | **PV: 1.9** (decreased) | SLC25A2 is decreased in PV (with SLC2A3 and SLC44A1 differentially expressed in ET). Solute membrane carrier proteins have been associated with venous thromboembolism in genome wide association studies and *in vivo* models<sup>97,98</sup>. |
| **RAB32** | **PV: 2.1** (increased)<br>**ET: 1.6** (increased) | RAB32 is increased in ET (along with mitochondrial membrane protein TOMM22). Mitochondria are recognized as key regulators of platelet procoagulant function<sup>99</sup>. Loss of mitochondrial protein RAB32 is associated with dense granule storage pool disease Hermansky-Pudlak syndrome<sup>100</sup>. |
| **PSMD11** | **PV: 2** (increased)<br>**ET: 1.7** (increased) | We show evidence of dysregulated protein degradation pathways with upregulation of PSMD11 along with differential expression of lysosomal proteins (SORT1 and ATP6V) and other proteasomal subunits. This reflects the work of other groups who have shown that protein quality-control pathways may be important in the pathogenesis of MPN and other prothrombotic diseases and represent novel therapeutic targets<sup>101-103</sup>. |

### Differential platelet proteome in MPNs identifies potential mediators of disease phenotype

To better decipher the functional and biological significance of the observed proteomic changes, we performed pathway-enrichment analysis and identified signaling pathways that are differentially activated between MPN subtypes and controls (**Figure 3**). Gene set enrichment analysis (GSEA, see Methods) found that MPN (stratified by subtypes; ET and PV) primarily induces pathways associated with the unfolded protein response. Impaired proteostasis is attributed^23,24^ to endoplasmic reticulum stress (note overexpression of candidate proteins HYOU, PDIA6, HSP90B1 in this data, **Figure 2B**). Moreover, among the most enriched gene sets, MPN pathology induces activation of oxidative phosphorylation (OXPHOS) and mTORC1 signaling pathways. Proliferation pathways also reveal significant enrichment with overexpression of c-MYC target proteins among patients with PV. Representative GSEA profiles are shown in **Figure 3** and the full list of gene sets are detailed in **Table S2A-B**. The MPN pathways exhibiting significant proteomic regulation by GSEA are consistent with our observations at the individual level for increased or decreased protein expression and mirror key signaling pathways identified in our prior platelet transcriptomic analysis^21^. These results are also consistent with independent studies in MPN granulocytes^25,26^ and hematopoietic stem and progenitor cells^27-29^ confirming their significance in MPN pathobiology and as potential therapeutic modalities^30^.

**Figure 3:**
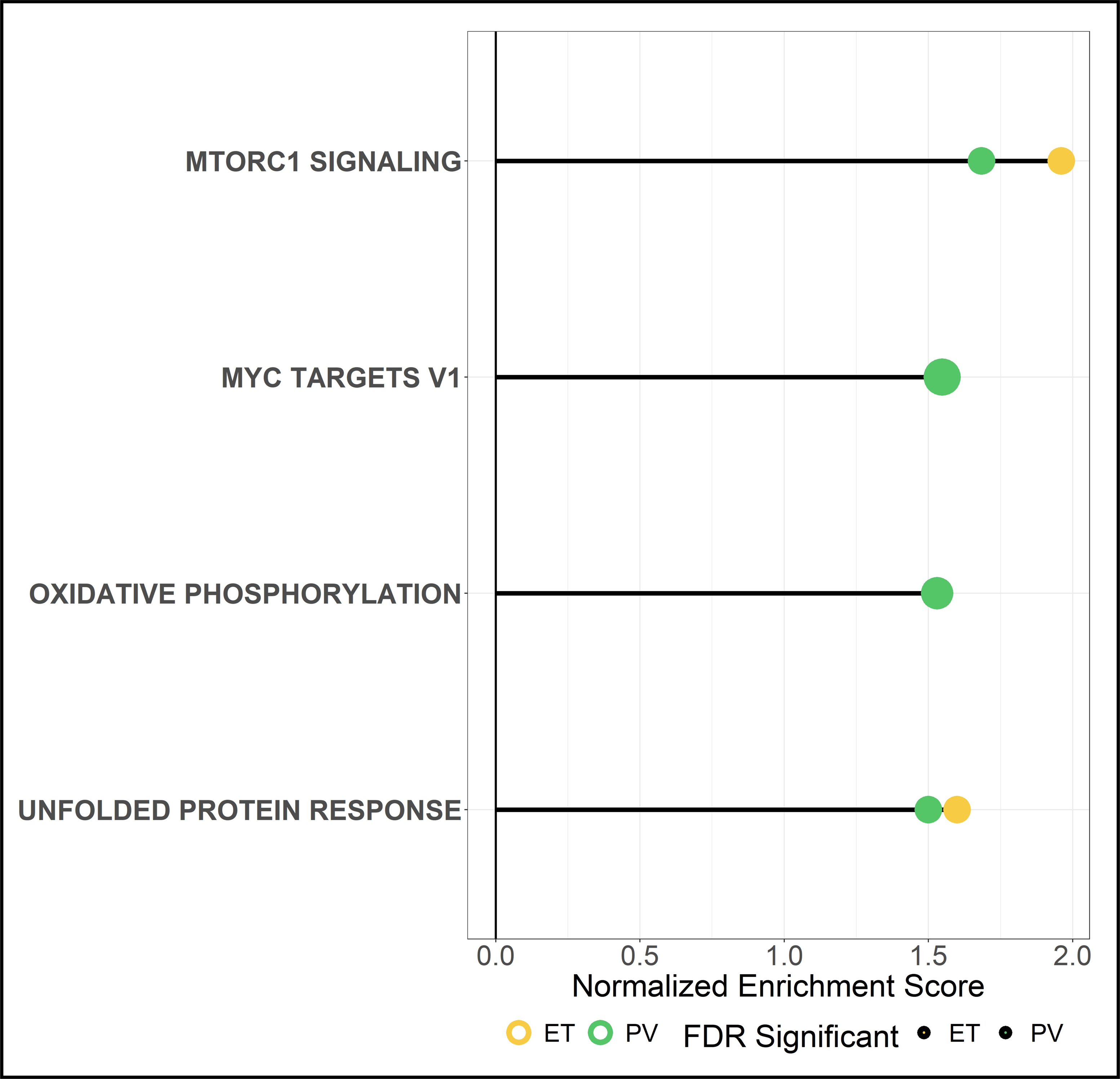
The platelet proteome in MPN identifies biological pathways recognised as crucial drivers of disease. **A,** Pathway-enrichment analysis of proteins with MPN subtype-specific expression (color indicated; yellow ET, and light green PV). Each point represents a pathway; the *x*-axis gives the normalized enrichment score, which reflects the degree to which each pathway is over-represented at the top of the ranked list of differentially expressed proteins, normalized to account for differences in gene set size and in correlations between gene sets and the expression data set. The y-axis lists the detail-level node of the most enriched pathways; solid lines mark GSEA-recommended^65^ Bonferroni-corrected statistical significance criterion of FDR < 0.25 for exploratory analyses.

### Shared platelet proteo-transcriptome by MPN subtype

Having defined differential protein signatures by MPN subtype, we proceeded to identify significant (FDR < 0.05) shared signatures between the current platelet proteomic data and our prior transcriptomic data^21^. We found 214 differentially expressed markers (significant ones alone across MPNs) at both transcript and protein levels – together translating to a moderate correlation (Spearman’s rho =0.44) between each other (**Figure 4**). In general, this is consistent with previous transcriptome-proteome comparative studies, including across species and matched cell-line experiments^31-33^.

**Figure 4:**
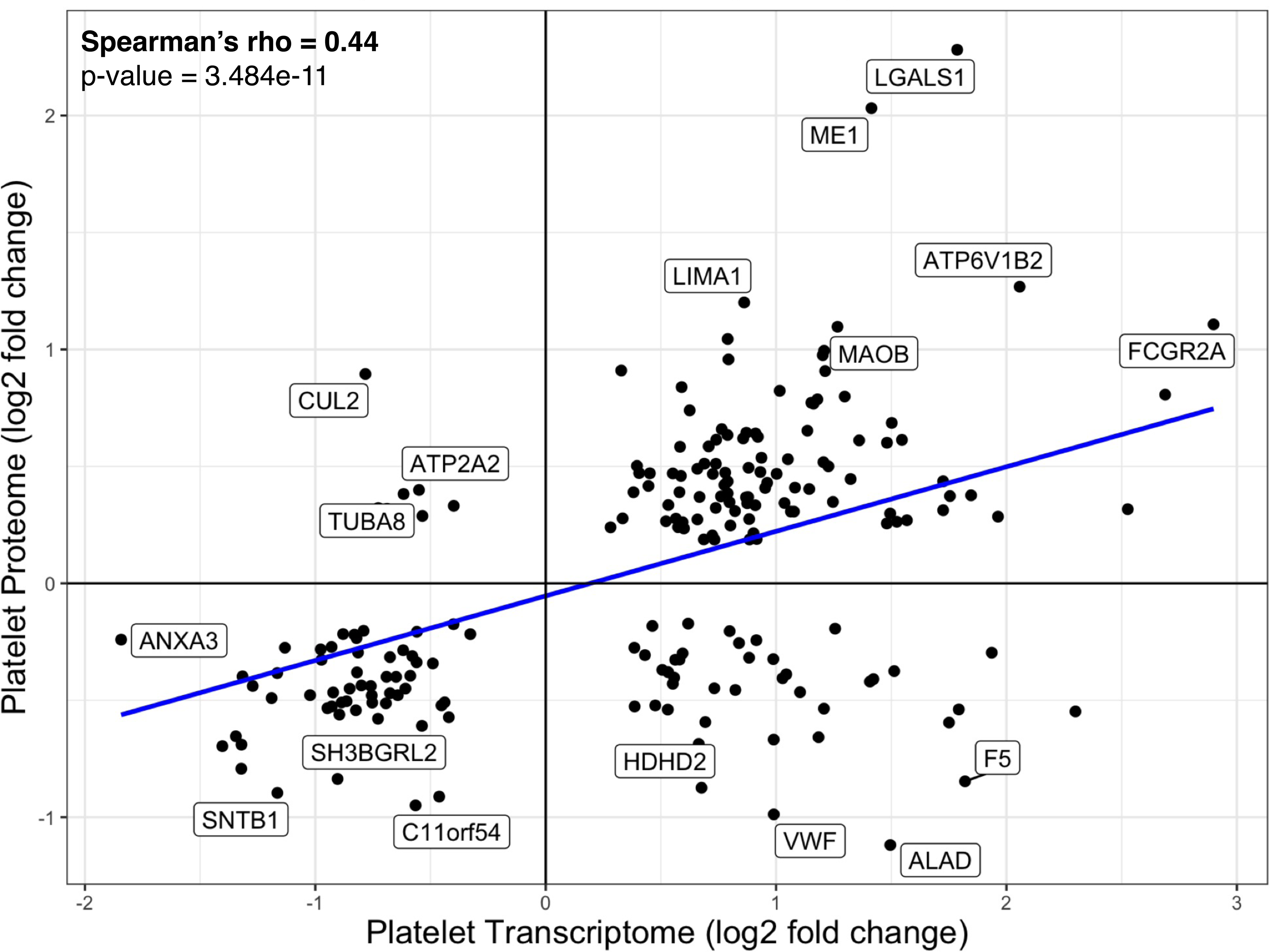
Correlation of differential expression between a shared set of significant genes and proteins. Spearman’s correlation showed moderate (Rho = 0.44, *p* = 3.484e-11) correlation in differential expression (log2FC) of genes from the MPN platelet transcriptomic cohort that were also significantly differentially expressed (FDR < 0.05) at the MPN platelet proteomic cohort.

Taking the overlap of the significantly differentially expressed proteins and genes (FDR < 0.05) that were common across MPN subtypes (i.e., present in both ET and PV specimens), we identified highly expressed features (n= 17) shared between the two independent MPN ‘omic’ cohorts. Unsupervised hierarchical clustering further classified this core MPN platelet proteo-transcriptomic overlap for ET and PV subtypes, in contrast with that of healthy donors. **Figure 5** demonstrates graded expression of 17 shared signatures between the MPN platelet proteome and transcriptome. Although MPN patients cluster into a group distinct from controls, we also note the overlay between PV and ET platelet signatures likely reflecting shared pathobiology. Most importantly, these signatures point to key molecular pathways previously identified in our MPN platelet transcriptome data and now validated at the protein level in the ET/PV platelet proteome (Figures 2, 3, and 5). These include, for instance impaired proteostasis and ER stress (known markers^34-36^ PSMD13, UNC45A, ATP6V1B2, NUCB1), oxidative stress and apoptosis^37-40^ (MAOB, PSAP, VDAC1, TIMP1) and lipid metabolism^41-43^ (ME1, DHCR7, FAH). Moreover, as shown earlier in **Figure 2B**, known markers of platelet activation, classical coagulation factors and mediators of immunothrombosis such as FcγRIIA, F5, VWF LGALS1, and LGALS3BP, are also confirmed.

**Figure 5:**
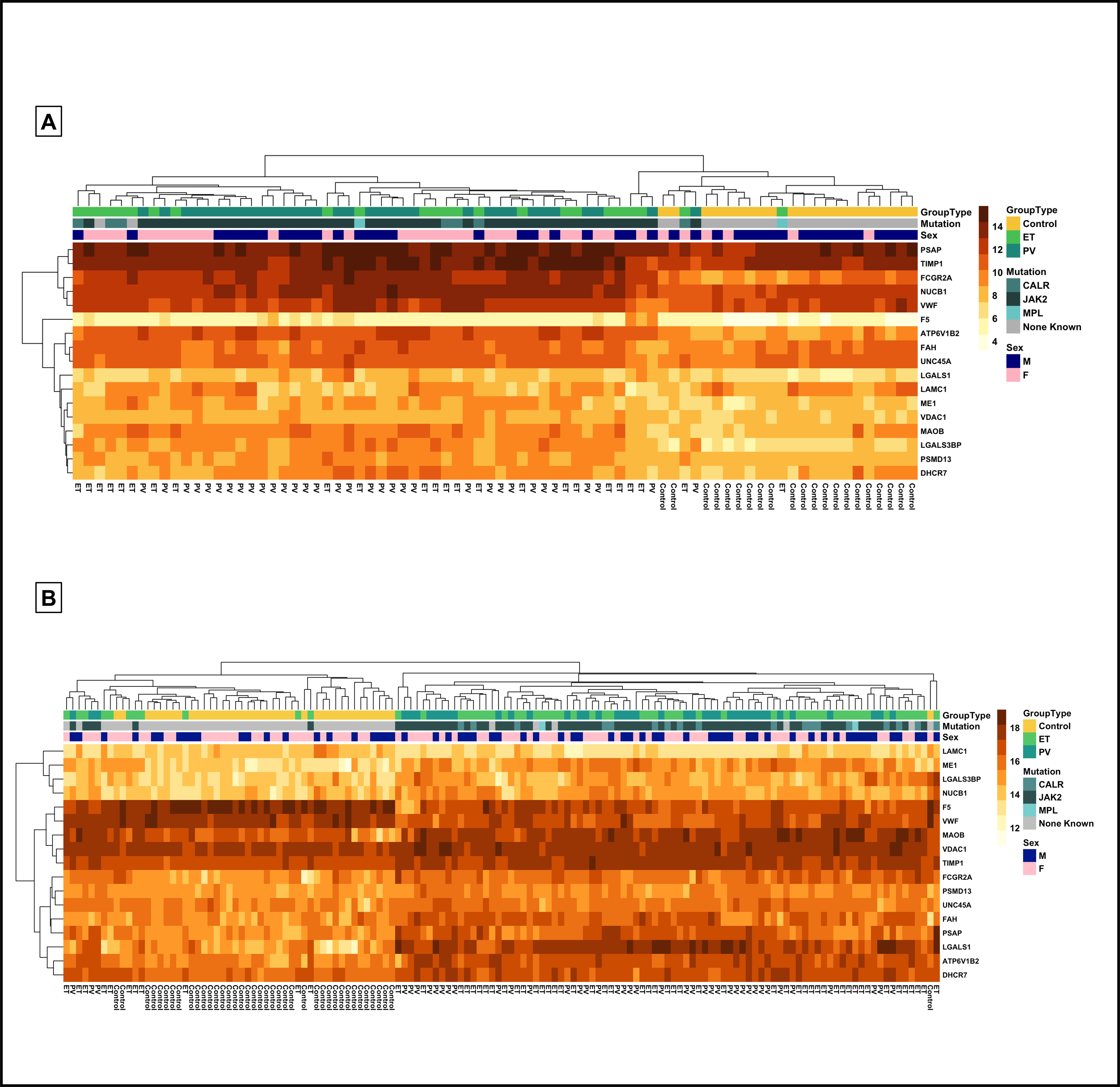
Highly expressed shared platelet proteo-transcriptome features in MPNs. **(A & B)** Taking the overlap of the significantly differentially expressed proteins and genes (FDR < 0.05) that were common across MPN subtypes (i.e., present in both ET and PV samples), we identified highly expressed features (n= 17) shared between the two independent cohorts. Hierarchically clustered heatmaps of the overlapping features between significantly expressed protein and genes (FDR <0.05) from controls versus MPN (present in both PV and ET). **A,** Transcriptomic expression from the Stanford single center cohort (n= 78). **B,** Proteomic expression from the international, multi-center collaborative (Ireland & Italy: n= 140). Colored annotation is provided to indicate MPN subtype, mutation status and sex. Rows indicate gradation in expression on a yellow (low) to orange (high) scale. Columns indicate sample type from controls, ET, and PV.

Thus together, this comprehensive proteo-transcriptomic analysis validating MPN platelet signatures not only between RNA and protein expression but also across two independent sites and clinical cohorts highlights additional underlying mechanisms beyond the MPN genetic landscape (and the known pivotal driver and non-driver mutations). The observed platelet molecular profile supports our hypothesis that platelets contribute to remodeling of the circulatory microenvironment which could lead to a self-reinforcing inflammatory milieu, possibly promoting disease progression and associated MPN vascular complications.

## Discussion

Here, we present a comprehensive proteo-transcriptomic study of blood platelet expression in patients with chronic progressive MPNs across international multi-site clinical cohorts (first of this kind to the best of our knowledge). Thrombocytosis is a cardinal feature of MPN, however platelet count in isolation is not predictive of clinical outcome^44^. Moreover, conventional anti-platelet therapy does not fully mitigate thrombotic risk and cytoreductive therapy has shown limited potential to alter natural history of the disease to date^45^. Recent data suggest^46^ that disease course may be modified with targeted treatments, however options remain limited, with incomplete understanding of why patients fail first line therapy with few available alternatives. One approach that is increasingly recognized as highly valuable in identifying novel therapeutic targets as well as molecular markers of disease (especially at a systems level) is to leverage pertinent multi-omic^47-49^ datasets.

In this study, we unravel the molecular phenotype of platelets, with a focus on the platelet proteome in the highly prothrombotic subtypes of MPN (PV and ET). Crucially, we validate this proteomic data with our prior MPN platelet transcriptomic data (ET, PV, and MF), confirming a strong possible role for platelet mechanisms in MPN pathobiology. Our data also expands on findings from other smaller or targeted studies, thus identifying the platelet proteo-transcriptome as potential mediators (and therefore, an invaluable biosource) of MPN proinflammatory, prothrombotic, and profibrotic processes^50-56^.

Our international collaborative effort demonstrates an example of addressing pertinent, global research questions (particularly for rare disorders such as MPNs). Importantly, this cross-institutional body of work emphasizes the value of standardized protocols in achieving larger cohort sizes and ensuring external validation toward high-quality and reproducible data of direct relevance to patient care^57,58^.

In conclusion, using label-free, untargeted platelet proteomic profiling, we discover key pathobiological mediators of thrombosis and proteostasis in MPN patients (focusing on ET and PV subtypes) from multiple recruitment sites; and validate the proteome with our prior transcriptomic data from an independent cohort.

We identify significant differential protein expression, that were also noted at the RNA level, thus strengthening biological insights, and revealing high priority candidate markers for functional evaluation in MPN studies. Our findings highlight the platelet molecular profile as a unique peripherally accessible window into MPN pathobiology. We also demonstrate the immense value of collaboration between groups with shared research goals and complementary expertise in increasing the feasibility of large-scale translational studies, particularly for rare disease cohorts.

## Limitations of the study

There are several potential limitations of this study. Firstly, the comparative multi-omic measurements were not in the same patients, but in separate MPN clinical cohorts at independent sites. Consequently, there were limitations on the extent of integrative multi-omic analyses possible. However, our results do align with a recent independent but smaller study with paired MPN platelet multi-omic analysis^50^. Of note, there is increasing acknowledgement that a ‘rectangular approach’ using untargeted, discovery methodologies from large, geographically independent cohorts of patients at both the discovery and validation phases of biomarker research is superior to traditional ‘triangular workflow’, where hypothesis-free technologies are used initially followed by targeted validation of a smaller candidate markers in a larger sample size^59^. Secondly, in line with our hypothesis focusing on platelets as prothrombotic mediators we have limited our investigation to PV and ET patient samples. However, our analysis revealed the heterogeneity of the platelet proteome and implicated platelets in a diverse range of biological pathways (and confirming signatures identified in our prior MPN platelet transcriptomic data). We recognize that a longitudinal study following patients for the development of thrombotic outcomes (at adequate power) would be critical and will enable the development of future risk prediction algorithms. Thirdly, data-dependent acquisition mass spectrometry was carried out in this study, wherein a subset of the most abundant ions reaching the mass spectrometer detector are individually fragmented. Incomplete protein coverage is a well described limitation of this technique, particularly for low abundant proteins, and we recognize that failure to detect RNA-protein correlates in our platelet samples does not suggest absence at the protein level. Finally, we recognize that future mechanistic studies are needed to substantiate results from this study focused on characterizing the MPN platelet proteo-transcriptome alone. Our ongoing efforts include proteomic analysis of platelets from prospectively recruited patients at the time of initial MPN diagnosis to identify candidates for mechanistic interrogation in treatment naive patients and to better understand the platelet molecular signature as a potential diagnostic, prognostic, or therapeutic target for patients with MPN.

## Methods

### Study recruitment and sample preparation for platelet proteomics

Ethical approval was granted from the Institutional Review Board (IRB) of the Mater Misericordiae University Hospital, Dublin, Ireland (IRB approval number 1/378/2241) and the Papa Giovanni XXIII Hospital, Bergamo, Italy (IRB approval number 1789/2013). Patients over the age of 18 with an established diagnosis of MPN (PV n= 41, ET n= 59) according to the World Health Organization diagnostic classification and a control group of healthy donors (n= 40) were invited to participate (2014-2022)^60^. Following informed consent, samples of whole blood collected in sodium citrate (0.105mol/L) were obtained by direct venipuncture. Platelets were isolated from platelet rich plasma (PRP) obtained by centrifugation of whole blood for 10 minutes at 400 *g* at room temperature (RT), according to a published method^61^. Briefly, PRP was diluted in 1:2 ratio with Krebs Ringer buffer (4mM KCl, 107 mM NaCl, 20 mM NaHCo_3_, 2mM Na_2_SO_4_, pH 5). After centrifugation at 1,000 *g* for 10 min at RT, the platelet pellet was resuspended in Krebs Ringer buffer supplemented with glucose (0.9 g/L, PH 6) and centrifuged a second time (1,000 *g*, 10 min, RT). This washing procedure was repeated twice, and the platelets were resuspended at a concentration of 1x 10^9^ platelets/mL in phosphate buffered saline (PBS) or PBS containing 1% Triton, snap frozen on dry ice and stored at -80 °C.

### Mass Spectrometry

Whole platelets were lysed in RIPA buffer (100 mM Tris pH 8.0, 300 mM NaCl, 2% Triton-X 100, 0.2% SDS, 1% sodium deoxycholate) with protease and phosphatase inhibitors (Roche). Samples were precipitated with 95% acetone overnight at -20 °C, centrifuged at 14,000 *g* at 4 °C for 10 minutes and the supernatant was removed. The protein pellet was resuspended in PBS and protein concentration was estimated by measuring absorbance at 280nm using a DS-11 spectrophotometer (DeNovix) as before^62,63^. Mass spectrometry sample preparation was performed using the commercially available PreOmics iST HT 192x kit (P.O.00067). In brief, 50 µg of protein was simultaneously lysed, reduced, and alkylated for 10 min at 95 °C and 1000 rpm, transferred to a cartridge and subsequently double-digested with LysC and trypsin at 37 °C and 500 rpm for 1 hour. Peptides were purified with repeated washes and eluted. Samples were evaporated at 45 °C and peptides resuspended in LC-load buffer. Digested peptides were loaded onto C18 trap columns (Evotip) and washed with 20 μL 0.1% formic acid (FA) followed by the addition of 100 μL storage solvent (0.1% FA). Differential proteomic signatures were established using label free liquid chromatography mass spectrometry (LC-MS) analyzed in a Bruker TimsTOF mass spectrometer connected to an EvoSep liquid chromatography system operated by the UCD Conway Proteomics Core facility.

Samples were loaded onto the Evosep One LC system and separated with an increasing acetonitrile gradient over 40 minutes at a flow rate of 250 nl/min at room temperature. The mass spectrometer was operated in positive ion mode with a capillary voltage of 1500V, dry gas flow of 3 l/min and a dry temperature of 180 °C. All data was acquired with the instrument operating in trapped ion mobility spectrometry (TIMS) mode. Trapped ions were selected for MS/MS using parallel accumulation serial fragmentation (PASEF).

Identified peptides from platelet samples were searched against a human FASTA (July, 2022) using MaxQuant (2.0.3.0) with specific parameters for trapped ion mobility spectra data dependent acquisition (TIMS DDA). In the main Andromeda search precursor, mass and fragment mass had an initial mass tolerance of 6 ppm and 20 ppm, respectively. The search included fixed modification of carbamidomethyl cysteine. Minimal peptide length was set to seven amino acids, and a maximum of two miscleavages was allowed. The false discovery rate (FDR) was set to 0.01 for peptide and protein identifications. The normalized protein intensity of each identified protein was used for label free quantitation (LFQ) as previously described^64^.

### Proteomic analyses

Statistical analysis of the LFQ intensities was performed using Perseus (version2.0.10) and R (version 4.3.1). Protein identifications were filtered to eliminate identifications from the reverse database, proteins only identified by site, and common contaminants. For downstream analysis, only proteins identified in at least 50% of samples in at least one group (control/ET/PV) were included. Missing values were imputed using the random forest method (Missforest package, R/Bioconductor). Data was log2-transformed and differential protein expression was established using the Limma software package within R/Bioconductor.

Differential protein expression was adjusted for batch, patient age, sex, and treatment (antiplatelet and cytoreductive therapy) as potential confounding variables within the linear model in Limma (design <-model.matrix(∼patientvar$Subtype+patientvar$Batch+patientvar$Age+patientvar$Sex+patientvar$ASAnum +patientvar$HYDnum). Controlling for multiple comparisons was performed using the Benjamini-Hochberg defined false discovery rate (FDR). Significant differential protein expression was pre-specified as proteins with an FDR < 0.05 and a fold change of 1.5 in MPN, as compared to healthy controls.

Continuous data were summarized as medians and IQRs and categorical data are presented as frequencies and percentages. To compare differences in clinical variables between healthy controls and MPN subtypes (ET and PV), we used violin and box plots and conducted Mann-Whitney *U* test for non-parametric data. For unsupervised clustering and visualization, we performed principal component analyses (identifying MPN subtypes by color). All analyses were performed using the R studio interface (version 2023.03.1+446).

### Proteomic quality control and validation analysis

To assess intra-donor platelet proteomic reproducibility, 6 patient samples were analyzed as technical replicates (5 in duplicate, 1 in triplicate). Pearson correlation coefficient (*r*) was performed on the log2 transformed LFQ-intensity of all proteins quantified (n= 1771) across technical replicate samples. To assess biological (inter-donor) variability in protein abundances, Pearson correlation coefficient (*r*) was performed on the log2 transformed LFQ-intensity of all proteins quantified (n= 1952) across biologic replicate samples (control n= 40; MPN n= 100).

### Pathway/Gene set enrichment analysis for differentially expressed (DE) proteins

Gene set enrichment analysis(GSEA)^65^, a well-established method for determining regulatory patterns in co-expressed genes, was performed on the entire DE protein set for each MPN subtype (PV & ET), using the Cancer Hallmarks gene sets from MSigDB^66^. The ‘GSEA Pre-ranked’ function was used with a metric score that combines fold change and adjusted p-value together for improved gene ranking. We used default settings with 10,000 gene set permutations to generate *p* and *q* values and compared MPN subtypes. In these analyses, to allow for a broad comparison, we assessed all transcripts that were differentially expressed according to FDR/adjusted p < 0.25 as recommended by the authors of GSEA ^65^.

### Transcriptomic analyses

For platelet transcriptomics, detailed methodology including platelet isolation, next generation RNA sequencing and platelet transcriptome analysis is outlined in the original publication^21^. Briefly, peripheral blood samples from 99 patients with MPN (ET n= 24, PV n= 33, MF n= 42) were obtained from the Stanford Cancer Institute Hematology Tissue Bank from December 2016-December 2019 and 21 healthy donors from the Stanford Blood Center. All blood samples were obtained with written informed patient or donor consent. Ethical approval was granted by the Stanford University Institutional Review Board (IRB approval #18329). RNA Sequencing was performed at the Stanford Genomics core. Platelet transcriptomic data were library-size-corrected, variance-stabilized, and log2-transformed using the R package DESeq2^67^.

### Comparative proteo-transcriptome analysis

Our published MPN platelet RNA sequencing dataset was used for comparative analysis with the MPN mass spectrometry proteomic data. Comparative analysis of these two independent ‘omic’ cohorts was established using direct feature overlap of significantly expressed genes and proteins (FDR <0.05). Shared candidate markers between the two independent MPN -omic cohorts were evaluated for further analyses. We generated heatmaps of these common highly significant genes and proteins using the pheatmap R package and its built-in functions for hierarchical cluster analysis on the sample-to-sample Euclidean distance matrix of the expression data for each independent dataset. Spearman’s rank correlation coefficient (rho) was used to correlate differential gene expression (log2fold change) of common significant (FDR < 0.05) genes and proteins between the independent MPN platelet proteomic and transcriptomic cohorts.

## Acknowledgments

This multi-institutional collaboration spearheaded by early-career investigators (S.K., B.K., and A.K.) would not be possible without the significant foundational efforts from senior investigators at each of our institutions. Specifically, we would like to thank Prof. Anna Falanga and her team at University of Milano-Bicocca and Papa Giovanni XXIII in Bergamo, Italy, Profs. Fionnuala Ni Ainle and Patricia Maguire at UCD Conway SPHERE Research Group, University College Dublin, Dublin, Ireland, and Profs. Holden Maecker and Jason Gotlib, and Ms. Cecelia Perkins at the Stanford MPN Translational Research Center and Stanford University.

S.K. received a Training Fellowship from the International Society of Thrombosis Haemostasis and the International Network of VENous Thromboembolism Clinical Research Networks (ISTH/INVENT) to complete this multi-site study with A.K. as host mentor at Stanford University. S.K. is additionally supported by a Wellcome Trust & Health Research Board Irish Clinical and Academic Training (ICAT) fellowship. This work was also funded by US National Institutes of Health grants 1K08HG010061-01A1 (NHGRI), 3UL1TR001085-04S1 (NCATS), and the MPN Research Foundation to A.K, and additional partial support from U01HG011762 (GREGoR, PI Professor Stephen B Montgomery at Stanford).

## Authorship Contributions

S. Kelliher, B. Kevane, and A. Krishnan conceived of the overall study and secured funding. S.K. designed the experimental plan with input from P.M., F.N.A., B.K. and A.K. S.K. and S.G. coordinated and performed the sample acquisition and processing according to established protocols. Z.S. and A.K. developed the code for computational analyses and A.K. and S.K. performed and interpreted the analyses. S. Madden and K.B. provided bioinformatic support. L.W. supported platelet protein preparation and C.S. operated mass spectrometry at the UCD Conway Proteomics Core Facility. Clinical samples, and annotation were kindly provided by A. Falanga and M.M., F.S. and S.G. at the Hospital Papa Giovanni XXIII, Bergamo, Italy. A. Fortune., S. Maung, M.F. provided clinical samples and annotation at the Mater Misericordiae University Hospital, Dublin, Ireland. S.K., and A.K. wrote and edited the manuscript. All authors critically reviewed and edited the manuscript. All authors approved the final manuscript.

## Data Sharing Statement

The mass spectrometry proteomics data have been deposited to the ProteomeXchange Consortium via the PRIDE^68^ partner repository with the dataset identifier PXD046324. RNA-sequencing data from this work (original FASQ files from paired-end sequencing of all 120 samples) is already deposited to the NIH genomic data repository dbGAP under public accession # PHS-0021-21. v1.P1.

## Supplementary Figures

**Figure S1:**
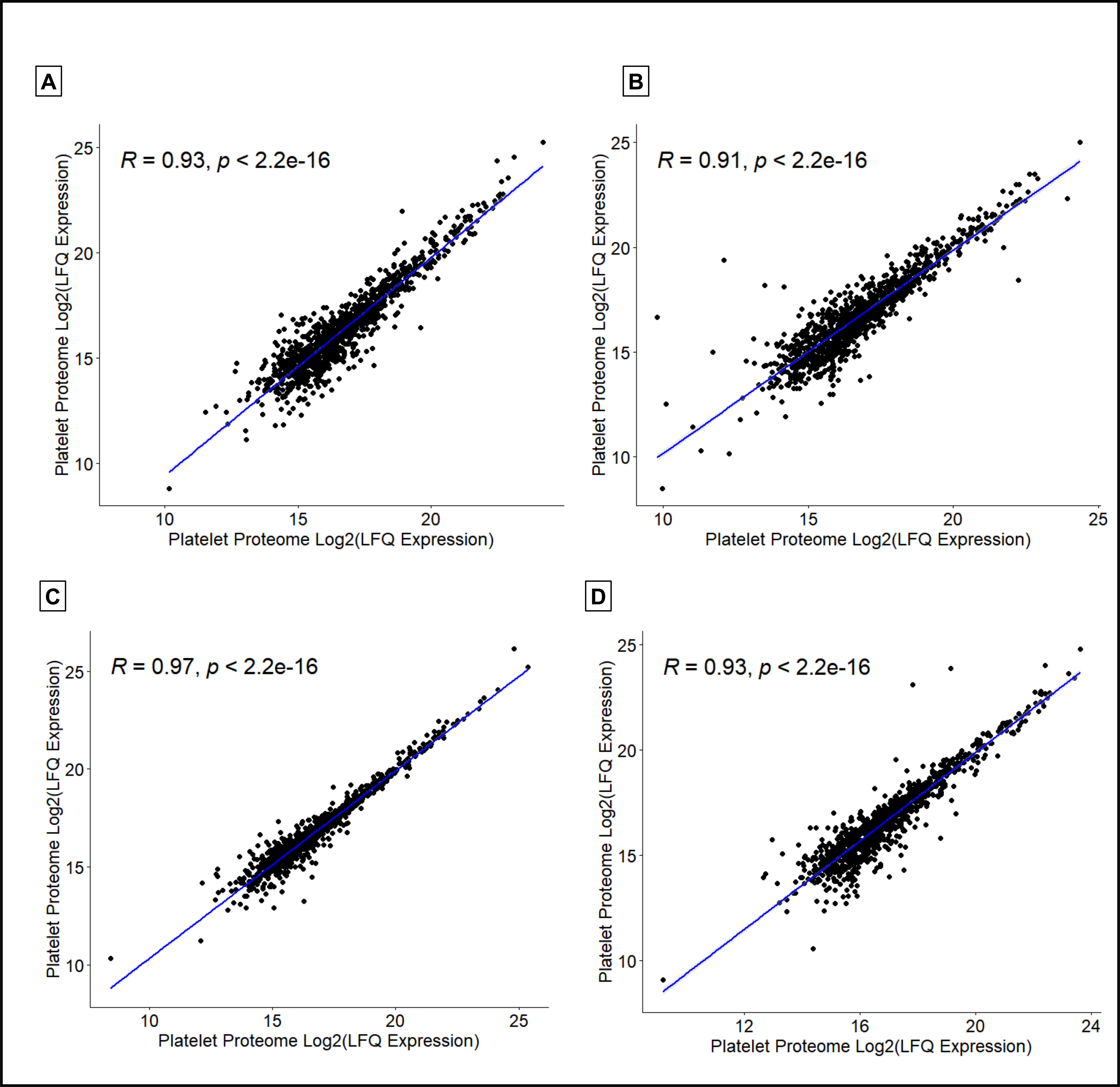
Strong correlation between platelet proteome technical replicates. Pearson correlation coefficients (*r)* from representative samples **(A-D**) demonstrate intra-donor reproducibility with strong correlation of log transformed LFQ intensities from technical replicates of the platelet proteome.

**Figure S2:**
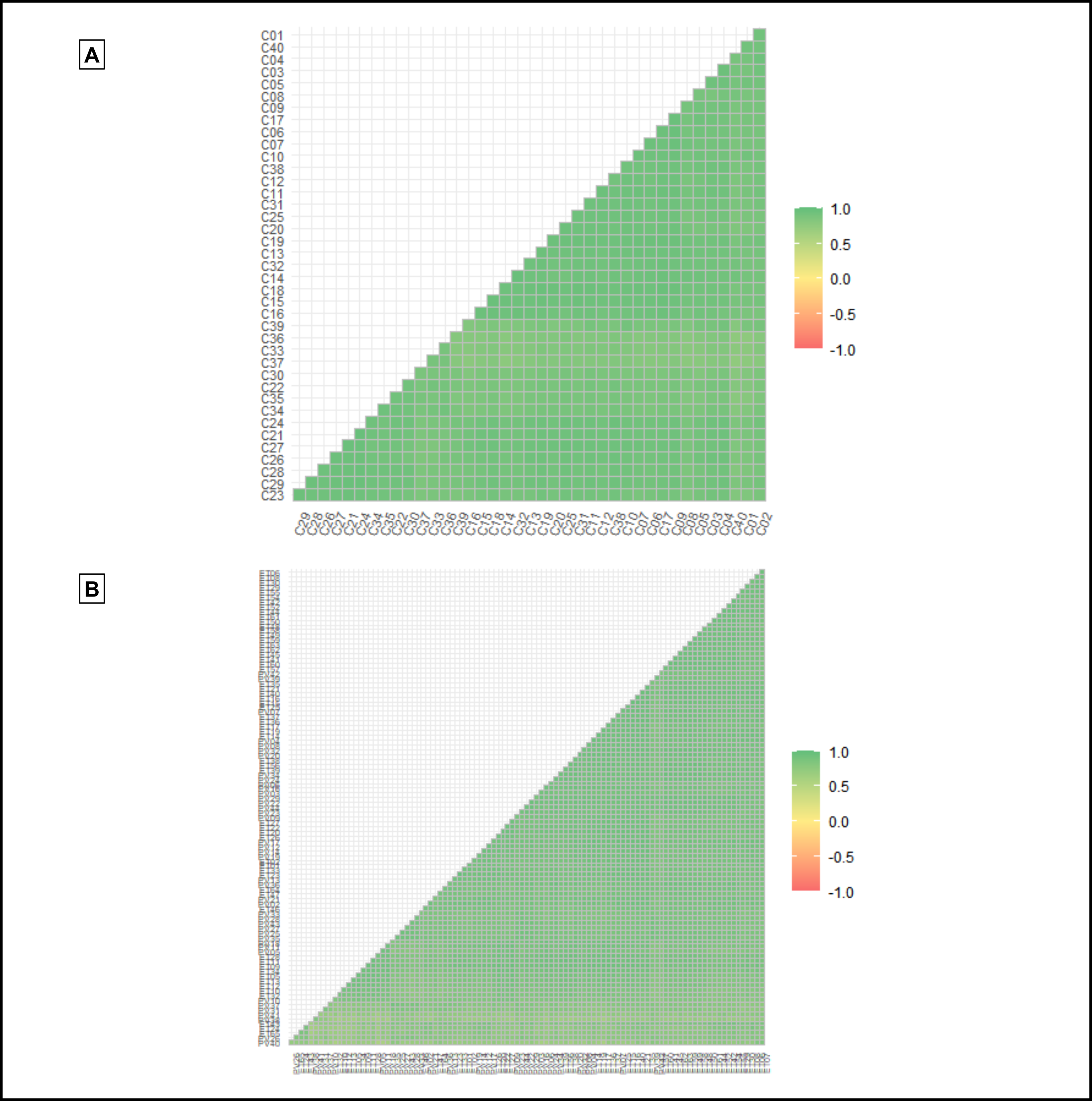
Strong correlation between platelet proteome biologic replicates. Correlation matrix of Pearson correlation coefficients (*r)* from biologic replicates show uniformly strong inter-donor reproducibility of the LFQ-proteomic analysis between biologic replicates from **(A)** controls (average *r* = 0.90 ± 0.04, min *r* = 0.75, max *r*= 0.97) and **(B)** MPN (average *r* = 0.88 ± 0.06, min *r* = 0.57, max *r* = 0.98) patient samples.

## Supplementary Tables

**Table S1:** 1952 proteins were quantified (LFQ intensity, see Methods) across PV, ET and control platelet lysate samples.

**Table S2:** 1315 proteins taken forward for downstream analysis. Proteins filtered to remove contaminants, proteins identified by site only, or in reverse. Proteins included were quantified across all groups (PV, ET and control) and were present in at least 50% of samples in at least one group (see Methods).

**Table S3:** Full list of 227 differentially expressed platelet proteins (Benjamini Hochberg false discovery rate <0.05) identified between ET and control samples.

**Table S4:** Full list of 166 differentially expressed platelet proteins (Benjamini Hochberg false discovery rate <0.05) were identified between PV and control samples.

**Table S5:** 1771 proteins quantified across technical replicate samples (n= 6).

**Table S6:** Full data for all molecular pathways identified in platelet proteome of ET patient cohorts.

**Table S7:** Full data for all molecular pathways identified in platelet proteome of PV patient cohorts.

**Table S8:** Correlation matrix with Pearson correlation coefficient *(r*) of log2 transformed LFQ intensity from biologic replicates of control (n= 40) samples.

**Table S9:** Correlation matrix with Pearson correlation coefficient (*r*) of log2 transformed LFQ intensity from biologic replicates of MPN (PV n= 41; ET n= 59) samples.

